# The structure of the Human Adenovirus 7 virus-like particles reveals that pentons and core-genome promote hexon-pIIIa interactions during capsid maturation

**DOI:** 10.1101/2025.09.22.677823

**Authors:** Kiyano Madoo, Ryan Mazboudi, Robert Alan Kuschner, Paul Gottlieb, Jose M. Galarza, Reza Khayat

**Author notes:** These authors contributed equally (Kiyano Madoo, Ryan Mazboudi).

## Abstract

Members of the *Adenoviridae* family routinely infect humans, exhibit significant genetic diversity, and are associated with a variety of illnesses. Types 4 and 7 frequently circulate in the United States and are major causes of respiratory disease. Infections can result in hospitalization and, in severe cases, death. Although a live wild-type-virus vaccine targeting these two types exists, its use is restricted to military personnel due to concerns about viral-shedding and the potential for genetic recombination. To overcome these limitations, we recently developed a virus-like particle (VLP) platform as an alternative vaccination strategy. These VLPs are stable, lack genomic material, and elicit a potent humoral immune response in mice, effectively neutralizing adenoviral infection. Here, we describe the cryo-EM structure of these VLPs at near atomic resolution. Structural insights are essential to ensure that the neutralizing antigens displayed on the VLPs accurately mimic those of the infectious virion, guide the design of particles with improved stability and efficacy, and enable engineering of VLPs with antigenic properties targeting multiple adenovirus types. The structure confirms that the key epitopes capable of eliciting neutralizing antibodies are appropriately displayed for antibody recognition. It also reveals previously unobserved interactions critical for capsid maturation. The penton base adopts multiple conformations when interacting with hexons, and its proper positioning is necessary to facilitate hexon-pIIIa interactions that stabilize the capsid. A comparison of the VLP with immature and mature adenovirus structures shows that core-genome packaging strengthens interaction between cementing proteins pIIIa and pVIII and the hexon shell.

**Significance Statement:** Adenoviruses are large and morphologically complex family of viruses that undergo a sophisticated process of assembly and maturation. These viruses are responsible for ocular, respiratory, and enteric infections, posing a particular risk to children and the immunocompromised. Despite their clinical significance, there are no adenovirus vaccines available to the public. Here we report the cryo-electron microscopy structure of human adenovirus 7 virus-like particles, which have shown strong potential as a vaccine candidate. Our analysis provides critical insights for the design of particles as vaccines or carriers of genetic therapeutics. The structure reveals how interaction between hexon and penton promote cement protein pIIIa for increased capsid interaction, and how core-genome packaging contributes to capsid maturation and enhanced capsid stability.

## Introduction

Human adenovirus (HAdV) was first isolated from human adenoidal tissue in 1953 by two independent research groups (1, 2). AdVs have been found to infect various vertebrates and are now classified into six genera: *Aviadenovirus* (birds), *Mastadenovirus* (mammals), *Ichtadenovirus* (fish), *Barthadenovirus* (reptiles, ruminants, marsupials, birds), Testadenovirus (turtles), and *Siadenovirus* (birds, tortoises, frogs) (3). These viruses are non-enveloped and possess an icosahedral capsid (pseudo-T=25) that encloses a linear double-stranded DNA (dsDNA) genome. Their genomes exhibit considerable variation, ranging from 26 to 48 kilobases (kb) in length and encoding approximately 30 to 40 genes. Similarly, their capsids range in size between 70 to 100 nm (4). There are more than 100 types of HAdV, which are classified into subgroups A to G based on serological and genomic analysis (5). Each subgroup exhibits specific tissue tropism, and infections can lead to a wide range of clinical manifestations that include hepatitis, myocarditis, gastroenteritis, conjunctivitis, bronchitis, pneumonia, and bronchiolitis (6). Outbreaks occur in settings with close contact, such as military barracks, dormitories, schools, and daycare centers (7-10). Although infections are typically self-limited, hospitalizations are common and fatalities have been reported of children, immunocompromised individuals, and healthy US military recruits (6, 11, 12).

Human adenovirus types E4 and B7 are among the most prevalent and problematic strains. Both spread rapidly through close contact, respiratory droplets, contaminated surfaces, and the fecaloral route. They are known to cause outbreaks of acute respiratory disease (ARD), ranging from mild, cold-like illness to bronchitis and pneumonia. Currently, there is no approved specific antiviral treatment for infection. Natural outbreaks of HAdV-4 and 7 have been particularly troublesome in military barracks. In the United States, approximately 80% of military recruits become infected, and 20% of those require hospitalization (13, 14). This high incidence of ARD led to the development of a live virus oral vaccine specifically for military personnel; however, it is not available to the public due to viral shedding and documented secondary transmission following administration (6, 15, 16). Moreover, the live virus vaccine conveys the risk of recombination, potentially generating novel subtypes (17). These concerns motivated us to develop an adenovirus virus-like particle (AdVLP) platform as an alternative f vaccination strategy (18). We successfully demonstrated that transient transfection of HEK 293 cells with genes encoding AdV-7 major capsid proteins (hexon, penton, and fiber), minor capsid proteins (pIIIa, pVI, pVIII, and IX), the chaperone L4 100K, and the accessory protein L1 52/55K leads to efficient AdVLPs production. These AdVLPs were morphologically comparable to wild-type AdV-7 virion, as confirmed by negative-stain electron microscopy. Furthermore, the AdVLPs remained stable for months when stored at 4°C and retained stability after lyophilization. Immunogenicity studies in mice demonstrated that the AdVLPs elicited a potent, sustained, and protective neutralizing immune response specific to AdV-7 (18). While the technique was initially demonstrated with AdVLP-7 production, multiple other types of AdVLPs have also been produced, including AdVLP-4, AdVLP-14, and AdVLP-55. The expression system represents a significant advancement not only for AdV vaccine production, but also as a next generation carrier for mRNA, a promising platform for novel gene therapy vectors. The VLP system is also a powerful tool for biophysical and structural studies aimed at elucidating the detailed mechanisms of AdV assembly, maturation, and stability. A key advantage of this expression system is its ability to accommodate targeted mutations, deletions, and insertions without being subject to the evolutionary selection pressures that affect infectious virion. Such pressure can lead to reversions, or the emergence of novel mutations that favor the production of more infectious progeny than the parental strain.

Here we report the first structure of an AdVLP. We determined the structure of AdVLP-7 using cryo-electron microscopy (cryo-EM) and identified key differences from previously published structures providing important new insights into AdV maturation. Notably, the AdVLP structure is unique in that it is unaffected by the influence of the core-genome. To our knowledge, no prior structural description of the AdV exists before its interaction with the core-genome. Consequently, it remained unclear whether, and how, core-genome packaging alters the conformations and interactions of proteins that define the AdV capsid. Using symmetry expansion, focused classification, and local refinement we show that AdVLP-7 particles are mosaics of icosahedral asymmetric units (iASU) - a phenomenon that, to our knowledge, has not been previously described. However, a survey of the deposited cryo-EM maps suggests that mosaic forms of AdV may have been observed in the past but were not thoroughly investigated. Specifically, we demonstrate that penton base (PB) adopts multiple conformations relative to the peripentonal hexon (PPH) and, in some cases, is absent from some iASU. To our knowledge, multiple hexon-penton interactions have not been previously described. We show that PB insertion into the capsid correlates with ordering of its N-terminus, along with enhanced interactions between PB-pIIIa and hexon-pIIIa, contributing to capsid stabilization. Finally, we visualize a trimeric fiber-knob connected to the PB. Together our structural data provides important insights into AdV capsid architecture prior to genome engagement.

## Results

### The mosaic AdVLP-7

We determined the structure of the AdVLP-7 at a nominal resolution of 4.2 Å using cryo-EM with single-particle analysis and icosahedral symmetry (Fig. 1). Diffuse density corresponding to the PB and the fiber suggests partial occupancy, deviation from icosahedral symmetry, and/or structural dynamics. Using the program OccuPy, we estimated the occupancies of the PB and fiber to be ∼60 and < 20%, respectively (19). Diffuse density is also observed beneath the PB -a similar feature reported by Condezo et al. and attributed to the L1 52/55K protein (Fig. 1) (20).

**Figure 1.**
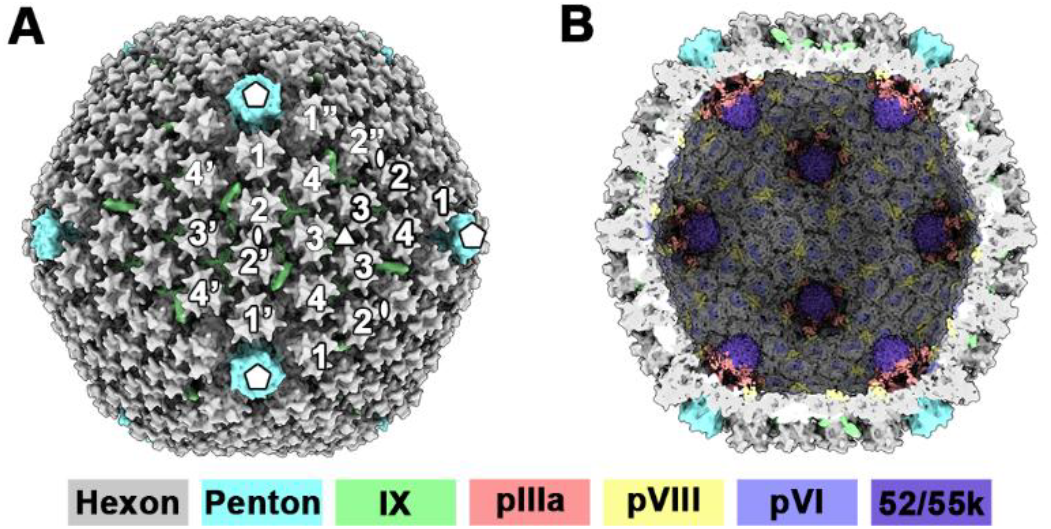
Schematic of the AdVLP-7 VLP. A) Surface representation of the low-pass filtered map showing the outer surface of the AdVLP in VIPER orientation. The icosahedral 5-, 3- and 2-fold axes of symmetry are labeled by a pentagon, equilateral triangle, and an ellipse, respectively. The twelve hexons defining a facet have been labeled 1-4 (light outline, medium outline, and dark outline), as historically defined. Hexons from adjacent facets are modified with prime (‘), and double-prime (“). B) Surface representation of the low-pass filtered map showing the inner surface of the AdVLP. Densities corresponding to proteins pIIIa, pVIII, pVI, and 52/55k are observed. The electron density representing each protein is color coded as shown at the bottom.

To determine whether the diffuse PB density results from limited occupancy or increased mobility, we used symmetry expansion and signal subtraction to isolate an iASU containing the associated PB. We then performed focused classification of the PB, identifying two distinct classes: one lacking (penton-less) and another possessing (penton-full) a PB (Fig. S1). Although the PB density in penton-full was significantly enhanced, it remained more diffuse than the neighboring PPH density. Therefore, we conducted an additional round of focused classification on the PB of the penton-full sub-particles, followed by local refinement. This analysis yielded five distinct classes (Table S1). Classes 1-3 exhibit strong density for the PB, but only diffuse density for the fiber (addressed below through additional classification). In contrast, Classes 4 and 5 show diffuse PB density (Fig. S1). Additional classification of classes 4 and 5 did not improve the PB density. The primary differences between classes 1-3 are rotation and translation of PB relative to the hexon shell, as well as the ordering of regions in pIIIa. These details are discussed below using atomic coordinates. To determine whether these classes originate from distinct AdVLP that conform to a single class, or from AdVLP representing mixtures of multiple classes (i.e., mosaics), we mapped each sub-particle back to the coordinates used for picking particles from the micrographs (i.e., the originating AdVLP). The analysis reveals that each AdVLP is composed of at least three different classes, with some incorporating all six classes. Therefore, we conclude that each of the imaged AdVLP is a mosaic of classes. The mosaic nature of the imaged particles may result from either the inherent nature of AdV, the expression protocol, the absence of a core-genome (affecting particle stability), the purification protocol, or destabilization during the wicking process used to prepare cryo-EM grids. Interestingly, both Barthadenovirus and ChAdOx1 demonstrate PB with an occupancy of ∼50% (Fig. S2).

### The penton base adopts multiple positions at the vertices

We generated atomic models for penton-less map and the three classes exhibiting PB density (Fig. 2, S3, S4, S5, Tables S1 and S2). For clarity, we refer to these four classes as C1 through C4, respectively (Fig. S1, Tables S1 and S2). The structures of hexon and penton were first reported by Athappilly et al., and Zubieta et al., respectively using X-ray crystallography (21, 22). The structures of AdVLP-7 hexon and penton are comparable to those reported for the infectious virion, with exceptions detailed below. Density corresponding to the hexon’s seven hypervariable regions (HVR) can be interpreted (Fig. S4). Structure comparison of the hexon protomers from the four classes yields root mean standard deviation (rmsd) values of less than 1.3 Å, indicating that the protomers are comparable between the four classes. However, overlaying the PPH protomers reveal hexon and penton conformational changes resulting from the insertion and movement of the penton (see below).

**Figure 2.**
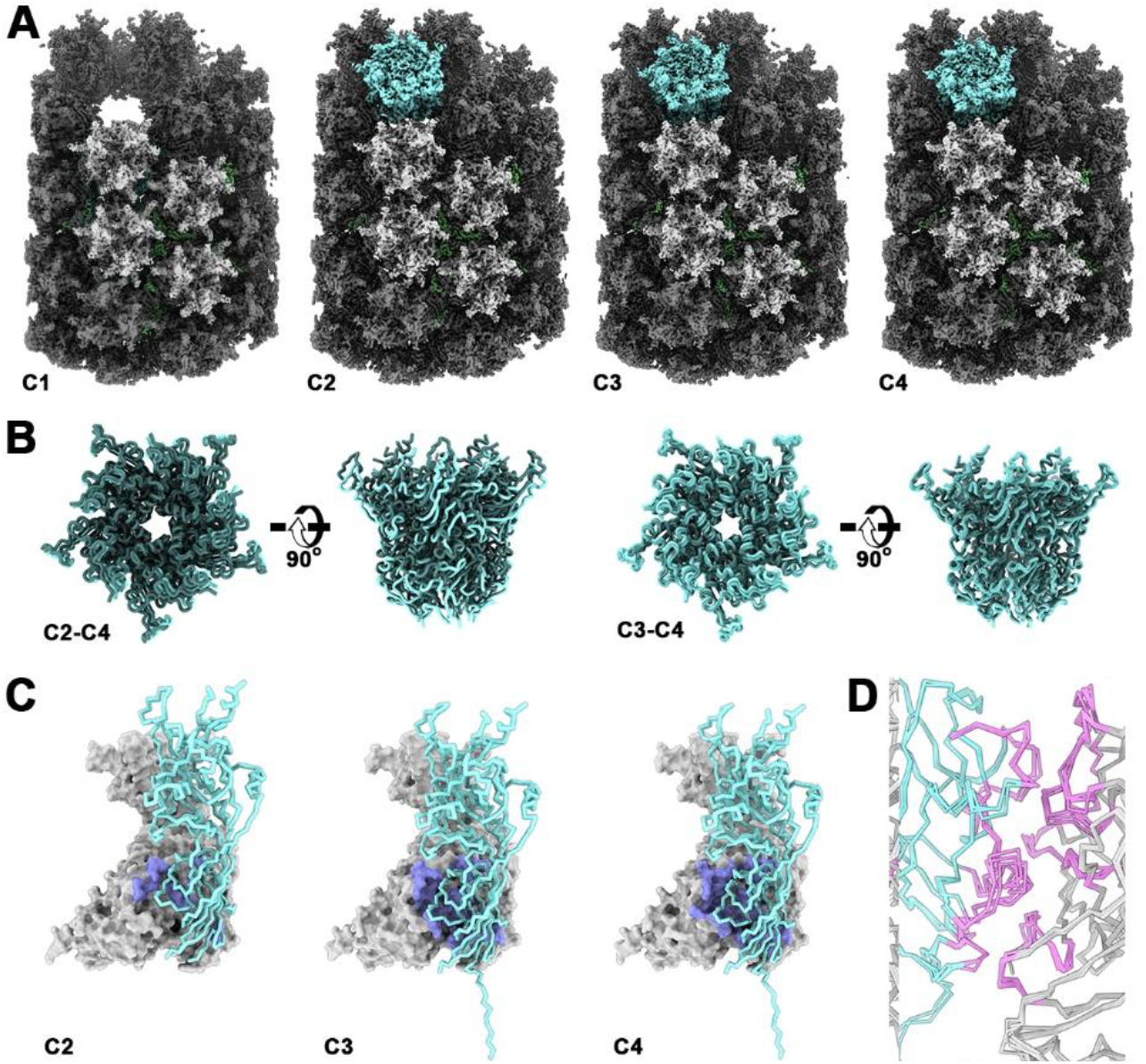
Pentons of the mosaic AdVLP-7. A) Surface representation of iASU classes (C1-C4) identified through symmetry expansion and focused classification. The multiple subunits are colored as Fig. 1. B) Comparison of the C2, C3 and C4 PB. The PB are aligned according to their associated PPH subunits then oriented along their 5-fold axis of symmetry. Left: Licorice representation comparing the C2 and C4. C2 is shown in dark cyan, and C4 in cyan. Right: Licorice representation comparing the C3 and C4. C3 is shown in medium dark cyan, and C4 in cyan. C) Surface and Cα trace representation identifying the hexon-penton binding interface in C2, C3, and C4 (left to right). The hexon is depicted as a grey surface, the penton binding interface is colored magenta, and the penton is shown as a Cα trace in cyan. D) Cα representation of hexon (grey) and penton (cyan) showing that both hexon and penton undergo conformational change in the different C1-C4 interfaces. The binding interface residues are colored magenta.

Interpretable density for C2 penton is observed for amino acids 52-306/350-543. Interpretable density for C3 and C4 penton is observed for amino acids 38-308/350-543 (Table S2). The region lacking density (307-349) is referred to as hypervariable region 2 (HVR2) and contains the integrin-binding loop (^329^RGD^331^) (Fig. S5). Overlaying the iASU coordinates for the three classes enables quantification of the penton positions relative to the PPH. Within the context of the AdVLP, the PB of C2 protrudes from the capsid shell, while the PB of C4 is inserted deepest into the capsid shell. The PB of C2 must shift by 7.8 Å (into the capsid) and rotate by 2.6° to overlay onto the PB of C4 (Fig. 2) The PB of C3 must shift by 1 Å (into the capsid) and rotate by 0.2° to overlay onto the PB of C4. The rmsd between the C3 and C4 PB before and after alignment are 1.7 Å and 0.6 Å, respectively. Insertion of the PB into the capsid is correlated with penton residues 38-51 becoming ordered and interacting with both the PPH and pIIIa in classes C3 and C4. A penton subunit contributes 900 Å^2^, 4200 Å^2^, and 4600 Å^2^ of buried surface area (BSA) per iASU for C2, C3 and C4, respectively (Table S3). The penton-pIIIa interaction is absent in C2, but accounts for 1500 Å^2^ of BSA for C3 and C4. Notably, the residues that define the hexon-penton interface in C2 are a subset of those in C3, and the residues in C3 are a subset of those in C4. Thus, as the PB becomes increasingly inserted into the hexon capsid shell, both the hexon-penton and penton-pIIIa interfaces expand.

### Plasticity at the hexon-penton interface

By comparing the structure of isolated PB with that of PB incorporated into the AdV-41 capsid, Rafie et al. were able to demonstrate that isolated PB undergoes conformational change upon insertion into the capsid (23). In this study, we describe a conformational change in the hexon upon PB insertion into the capsid. To investigate this, we generated two root mean square fluctuation (rmsf) plots: one for the C2-C4 pentons and another for the interacting PPH protomers. In each plot, we identify regions exhibiting significant fluctuations and then map these on to the hexon-penton interface (Fig. 2). Penton regions located at the PPH-PB interface show high rmsf values because they undergo conformational changes necessary to interact with different hexon interfaces. The same is true in reverse for the hexons. In the pentons, these fluctuating regions include residues 74-80, 97-109, and 426-429 (Table S4). Contiguous hexon regions at the PPH-PB interface that exhibit high rmsf values include amino acids 75-78, 331-334, 340-345, 676-679, and 927-936. (Table S4). Taken together with the findings of Rafie et al., our results demonstrate that PPH and PB mutually influence each other’s conformation.

### Insertion of the penton increases the hexon-pIIIa and penton-pIIIa interface

The precursor of IIIa (pIIIa), a substrate of adenovirus protease (AVP), remains undigested in the AdVLP-7 (18). A single pIIIa per iASU is located on the interior surface of the capsid shell, where it contacts the PB, the PPH, hexons 2 and 3, and pVIII. Residues 28-104 (C1-C4) form two antiparallel helix-loop-helix motifs (*hlh1* and *hlh2*) positioned directly beneath the PPH (Fig. 3). In C2, an additional four residues (105-108) can be modeled. Residues 108-134 (C3 and C4) form a connector helix (*ch*) extending from PB towards pVIII. Residues 137-218 and 226-266 constitute an α/β domain situated beneath and interacting with pVIII (C3 and C4). The density corresponding to residues 105-266 progressively improves from C1 to C4, as PB concurrently inserts deeper into the capsid (C2-C4) and strengthens its interaction with both the PPH and pIIIa. Thus, these structural changes reflect a dynamic pIIIa (C1-C2) that becomes increasingly stabilized upon interaction with PB in C3-C4. The total BSA involving a pIIIa subunit is 2,000, 2,100, 5,700, and 5,700 Å^2^ for C1 through C4, respectively (Table S3).

**Figure 3.**
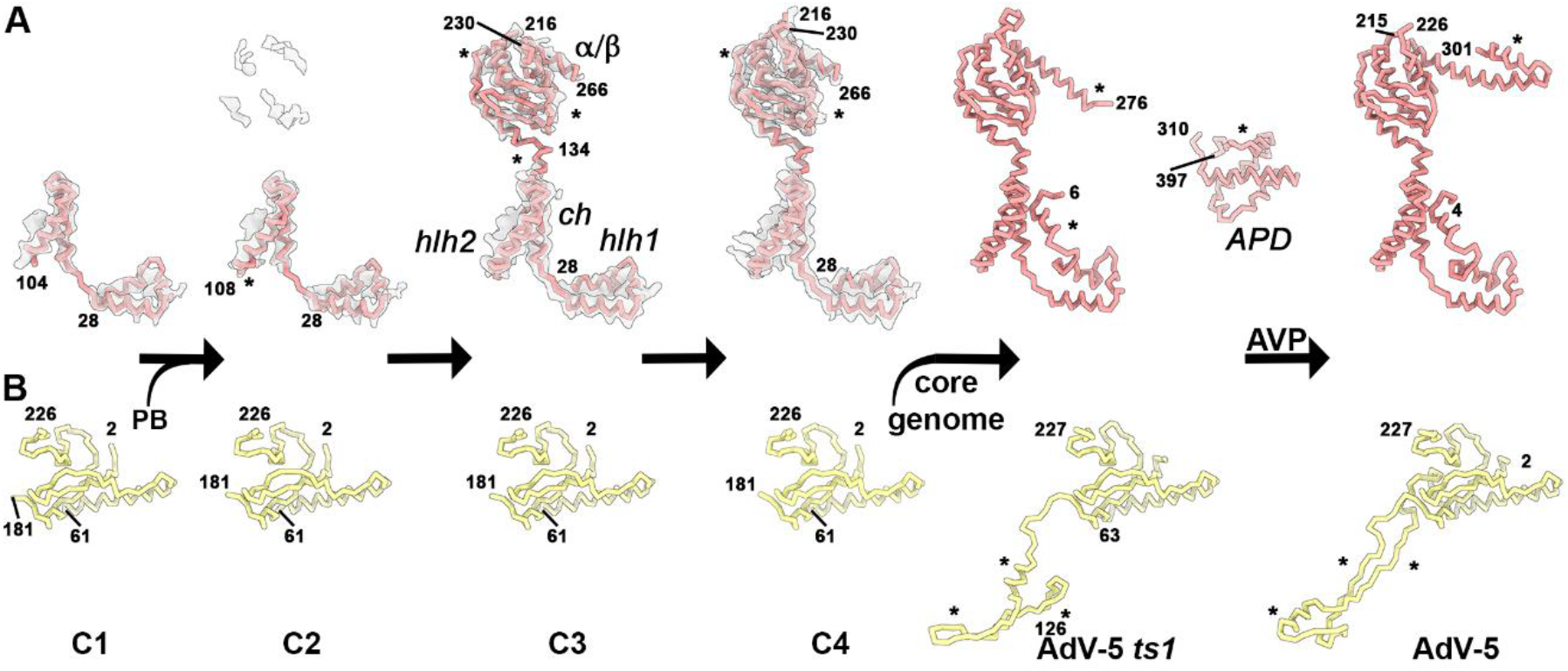
Conformational changes in AdV proteins pIIIa and pVIII induced by PB insertion, core-genome packaging, and AVP mediated proteolysis. Protomers colored as described in Fig. 1. The six structures analyzed included C1-C4 (this work), the AdV5 *ts1* variant (published), and mature AdV5 (published). Asterisks identify regions of structural difference between each panel and the one preceding it. A) Cα trace of proteins pIII and IIIa. In the absence of PB (C1), only the first two helix-loop-helix motifs (*hlh1 and hlh2*) are visible (C1). Insertion of PB into the capsid leads to conformational change of *hlh2* (C2). Further PB insertion and direct contact between PB and pIIIa, drives conformational change of the connector helix (*ch*) and the α/β domain (C3 and C4). Core genome packaging alters the N-terminus to nestle against *hlh1* and *hlh2*, extends the helix following the α/β domain, and results in icosahedral ordering of the appendage domain (APD). Subsequent proteolysis by AVP restructures the helix following the α/β domain and leads to icosahedral disordering of the APD. B) Cα trace of the pVIII and VIII subunits positioned beneath the PPH. PB insertion does not affect pVIII structure; however, core-genome packaging significantly stabilizes its interaction with the capsid shell (compared C1-C4 with AdV5 *ts1*), AVP-mediated proteolysis induces additional conformational change of IIIa. The pVIII subunit adjacent to the icosahedral 3-fold axes exhibits similar behavior.

The precursor of protein VIII (pVIII), also a substrate of AVP and remains undigested in AdVLP-7. There are two copies per iASU located in the interior surface of the capsid shell (chains U and V). One copy (pVIII^U^) is positioned beneath and interacts with two neighboring PPH, two neighboring pIIIa subunits, and hexon 4. Density is observed for amino acids 2-61 and 181-226 in C1-C4. The structure is predominantly composed of loops, with two α-helices, and a two stranded β-sheet. The second copy (pVIII^V^) also lies beneath and is associated with hexons 2, 3, and 4 of the same iASU, as well as hexon 3 of a neighboring iASU. Density is observed for amino acids 2-16 and 181-227 for all four classes. The two pVIII subunits interact with both intra- and inter-facet proteins to enhance capsid stability. Together, they contribute BSA of 15,500, 16,600, 18,200, and 19,200 Å^2^ to the C1-C4 interfaces, respectively. The increased interfaces in C3 and C4 are attributed to interaction with the ordered C-terminus of pIIIa (Table S2 and S3).

### Packaging of the core genome increases the hexon-pIIIa, penton-pIIIa, hexon-pVIII, and penton-pVIII interface

The hexon, penton, pIIIa, pVIII, IX, VI, and Fiber of AdV-7 and AdV-5 share 80, 74, 78, 79, 50, 66 and 30% sequence identity, respectively. The structures of hexon, penton, pIIIa, and pVIII overlay with rmsd values below 1.6 Å, suggesting that both viruses undergo comparable conformational adjustments during AdV maturation, which involves cleavage of precursor proteins by the DNA-dependent AVP. The structure of the AdV-5 *ts1* describes an intermediate where the core-genome has been packaged but AVP remains inactive and unable to process its substrates. In contrast, the mature AdV-5 describes the structural result of processing by an activated AVP (24, 25). Thus, the C1-C4 structures of AdVLP-7 provide insight into AdV capsid maturation prior to core-genome packaging, and AVP processing (Fig 3, 4). When compared to C1-C4, both pIIIa of AdV-5 *ts1* and IIIa of AdV-5 have over 20 additional amino acids modeled at their N-termini. In both structures, this region adopts a loop-helix-loop conformation that interacts with PPH. In AdV-5 *ts1*, the pIIIa structure extends from the C-terminus of C4 into a purely helical domain named the appendage domain (APD) which lies beneath the PPH, hexon 4, an adjacent PPH, and a neighboring hexon 2 (25). In AdV-5, IIIa extends from the C-terminus of C4 into a helix followed by a loop-helix structure, with the terminal helix interacting with hexon 4. Each pIIIa and IIIa subunit in AdV-5 contributes approximately ∼11,000 Å^2^ and ∼9,800 Å^2^ of BSA, respectively, to stabilize the immature and mature capsids (Table S3). The increased interface area arises from enhanced interactions between hexon and pIIIa, pIIIa and pVIII (or IIIa and VIII), and ordering of APD. Compared to AdVLP-7 pVIII, the pVIII of AdV-5 *ts1* and the VIII of mature AdV-5 chains contain over 50 additional amino acids that interact with both intra- and inter-facet hexons (Fig. 3, 4). These chains contribute approximately 27,600 Å^2^ and 27,900 Å^2^ of BSA, respectively, stabilizing the AdV-5 capsid.

**Figure 4.**
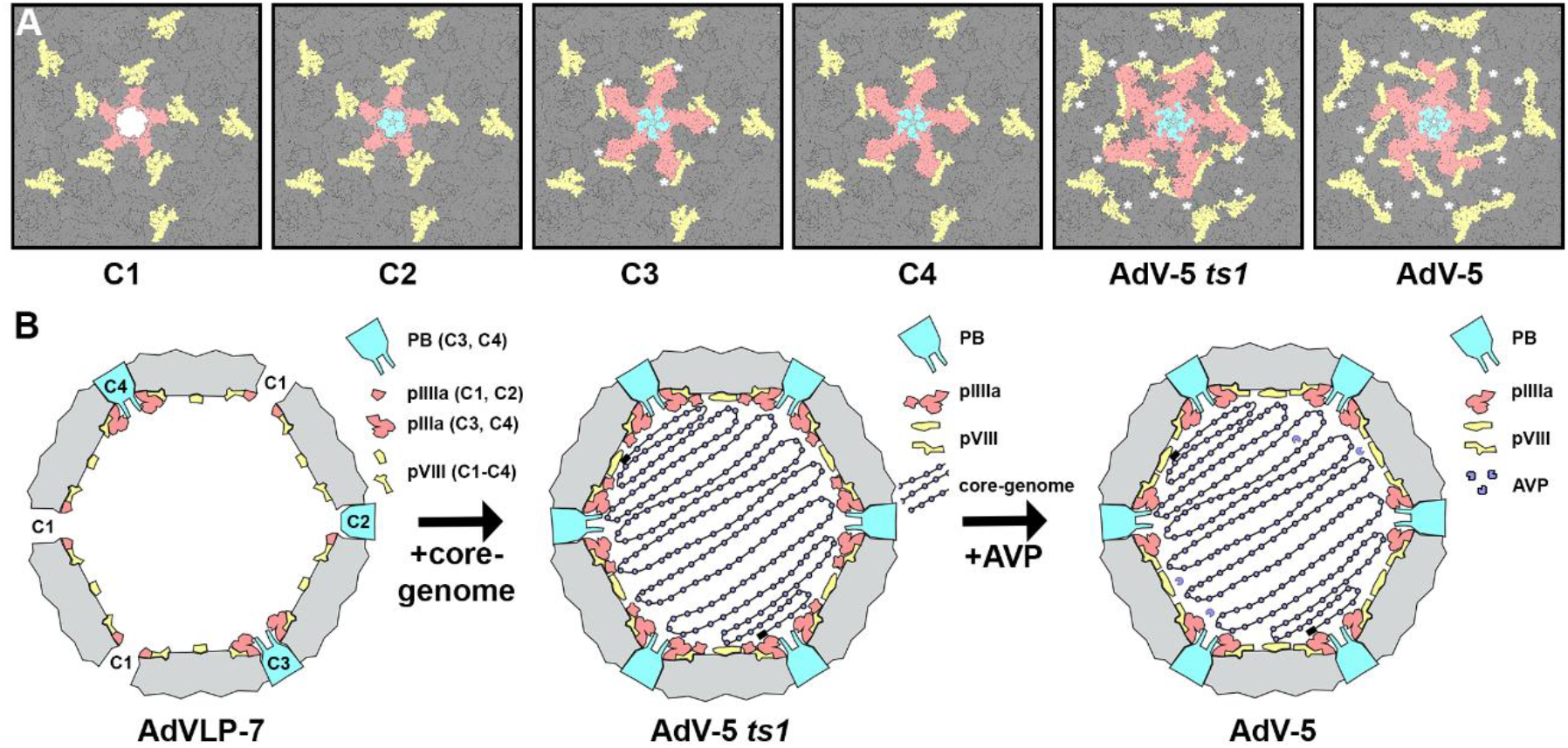
The core-genome anchors proteins for increased capsid interactions. A) Goodsell figures showing the icosahedral 5-fold axis of symmetry for C1-C4, AdV5 *ts1*, and AdV5. Subunits are colored according to Fig. 1. Insertion of PB promote hexon-pIIIa interaction -compare C1 to C4. Asterisks identify regions of structural difference between each panel and the one preceding it Packaging of core-genome promotes hexon-pVIII interaction -compare C4 to AdV5 *ts1*. Proteolysis of pIIIa by AVP alters hexon-IIIa interaction. -compare AdV5 *ts1* to AdV5. B) Cartoon representations of the mosaic, AdV5 *ts1*, and AdV5 capsids.

The primary difference between the AdV-5 *ts1* and AdVLP-7 is the presence of core-genome. We propose that in the absence of the core-genome, as in AdVLP-7, the pIIIa APD remains dynamic and is not observed. In contrast, packaging of the core-genome -whether through direct or indirect interaction-leads to stabilization and ordering of this domain, as seen in AdV-5 *ts1*. Similarly, we propose that the core-genome of AdV5-structures is responsible for positioning the 50 amino acids of pVIII to enhance capsid interaction. Proteolysis of pIIIa by AVP induces a conformational change of IIIa and leads to disordering of the APD (Fig. 4).

### The pentameric PB and the trimeric fiber knob

The fiber protein forms a homotrimer attached to PB, extending into the surrounding solvent (26). Structurally, the fiber resembles a globular knob positioned at the end of a shaft. The density corresponding to the fiber in the 4.2 Å icosahedral map appears diffuse likely due to limited protein occupancy and well-known flexibility of the shaft. Focused classification, described above, revealed diffuse fiber density in all classes containing the PB, indicating, therefore, that fiber association with the PB is independent of the PB’s position within the capsid (Fig. S1). We returned to the penton-full sub-particles, generated a cylinder mask using EMAN2.99, applied the mask to the PB, performed signal subtraction with Relion, and conducted focused classification to generate five distinct classes (Fig. S1). Only a single class displays density corresponding to the fiber, revealing a trimeric knob and a shaft with five repeating units (Fig. S1). To our knowledge, this is the first reported instance of trimeric density for the knob attached to PB. High-resolution refinement did not improve the resolution or quality of maps.

## Discussion

The only FDA approved effective vaccine for AdV is the live unattenuated AdV-4 and AdV-7 virus vaccine administered by the US military to all new basic recruits and other at-risk populations. However, this vaccine is not approved for public use. The AdV7-VLP platform represents a highly promising alternative to the live virus vaccine. Unlike the live virus, the AdVLP-7 is non-replicating and lacks genomic material, making it safer alternative. It can be efficiently expressed and purified in large quantities and induces high levels of protective neutralizing antibody in mice (18). Additionally, the VLP remains stable for weeks at 4 °C, can be lyophilized, is amenable to production of a range of AdV types, and can be engineered to display antigens from other viruses (18). Here, we provide near-atomic resolution data to confirm that no significant differences exist between AdVLPs and virions. The structure facilitates the engineering of AdVLP with enhanced assembly, stability, and efficacy. Our investigation has also demonstrated that the VLP platform offers a unique opportunity to investigate AdV assembly and maturation (Fig. 4). The structural information aligns with experimental evidence showing that PB dissociates from the capsid within the endosomes, promoting cellular entry and capsid disassembly (27). Moreover, atomic force microscopy (AFM) studies have shown that capsids subjected to external pressure respond by promoting PB dissociation, followed by capsid disassembly and genome release (28).

AdV particles are large (150 MDa), morphologically complex structures composed of many proteins. They undergo a sophisticated assembly process followed by a maturation step that is essential for infectivity (26). Two primary models have been proposed to explain how proteinaceous capsids enclose nucleic acids: the sequential and the concomitant pathways. The sequential pathway, extensively characterized for tailed bacteriophage, involves the assembly of empty capsids followed by genome packaging via a molecular motor powered by ATP binding and hydrolysis. The concomitant pathway entails the simultaneous assembly of the proteinaceous shell and encapsidation of the viral genome (29). Data supporting the sequential model comes from immunogold labeling experiments, which localize a complex composed of a potential terminase (IVa2) and a portal (33k) to a single vertex of the capsid. Metabolic labeling, Western blots, and quantitative mass spectroscopy estimate that each capsid contains 6-8 molecules of IVa2 (30). Structure prediction suggests that IVa2 functions as an ATPase, and mutations of its Walker A and B motifs abolish the formation of infectious virion (31). Negative stained electron microscopy images visualize the 33k protein to form ring-like oligomers with a central channel (32). The observation that AdVLP can assemble in the absence of AdV genome, IVA2 and 33K supports the sequential assembly model. The AdVLP-7 structure, which lacks some PB, is consistent with data showing that IVa2 and 33k are recruited to a unique vertex to package the viral genome into a preformed capsid shell. Evidence supporting the concomitant pathway includes electron microscopy images of freeze-substituted thin sections derived from HEK293 cells infected with an AdV-5 mutant exhibiting a delayed packaging phenotype. These images reveal partially formed capsids enclosing nucleic acid near the inner periphery of the nucleus. The authors propose that interaction between L1 52/55kD protein and viral DNA initiates capsid protein recruitment, leading to the formation of genome containing capsids. Disruption of this pathway, they argue, results in the accumulation of distinct pools of empty capsids and condensed DNA (33). Our system demonstrates that capsid assembly requires, at most, nine proteins (18). The ability of these proteins to self-assemble into AdVLPs clearly demonstrates that the viral genome is not required for capsid formation, and that successful virion assembly via the concomitant pathway likely depends on a tightly regulated process that ensures all components are correctly positioned to enable proper genome encapsidation. Unregulated expression of these proteins could lead to the accumulation of pools of empty capsids, consuming significant cellular energy and impairing efficient virion biogenesis.

## Materials and Methods

### AdVLP expression and purification

Expression and purification of AdVLP-7 was previously described (18). Briefly, AdV-7 (Genbank entry: AY594255) DNA sequences for hexon, penton, fiber, pIIIa, pVI, pVIII, IX, L4-100k, and L1-52/55k were codon optimized, synthesized and cloned into transfer vectors by Blue Heron Biotech (Bothell, WA USA). The genes were then sub-cloned into four expression plasmids: **i**. pcDNA3.4-hexon-IRES-100k (pHexon-100k), **ii**. pcDNA3.4-penton-IRES-fiber (pPenton-Fiber), **iii**. pcDNA3.4-CMV-VIII-IRES-VI-CMV-IX-IRES-IIIa (pVIII-VI-IX-IIIa), and **iv**. pcDNA3.4-L1-52/55k (p52/55k). Cloning was confirmed with restriction enzyme digestion and sequencing. The four plasmids (in a 2:1:1:1 ratio) were transfected into HEK-293-cells by mixing with PEI Max® (Polysciences, Warrington, PA, USA, 24765) at a 4:1 (PEI:total DNA) ratio in a volume of EX-CELL® CD HEK293 Viral Vector medium (Sigma Aldrich, St. Louis, MO, USA, 14385C) equal to 5% of the total culture volume. HEK-293 cells were cultured in suspension using EX-CELL® CD HEK293 Viral Vector medium (Sigma Aldrich, St. Louis, MO, USA, 14385C) supplemented with 5 mM L-glutamine. Cultures were maintained in a humidified incubator at 37 °C with 5% CO_2_. HEK-293 cells were while agitated at 125 rpm. Valproic acid, a histone deactylase inhibitor shown previously to increase protein production of transiently expressed genes, was added to cells 24 hours after transfection at a final concentration of 3.75 mM. At 72 hours post-transfection, HEK-293 cells were collected via centrifugation at 500xg for 10 min, and suspended in with 50 mM NaCl, 1 mM MgCl_2_, and 1× Halt™ protease and phosphatase inhibitors (Thermo Scientific, Waltham, MA, USA, 78440). The cells were subjected to three freeze-thaw cycles for lysis, centrifuged at 10,000 ×g for 20 minutes at 4 °C, and the supernatants concentrated tenfold using tangential flow filtration with a Pellicon® XL50 system equipped with Biomax® 300 kDa membrane (Millipore Sigma, Burlington, MA, USA, PXB300C50). Supernatants were layered onto a two-step cesium chloride (CsCl) gradient (1.41 g/mL and 1.26 g/mL) and subject to ultracentrifugation in a SW28 rotor at 10 °C for 2 hours. AdVLP-7 samples were visualized as a single band. Each gradient tube was punctured using an 18-gauge needle to collect fractions corresponding to the band of interest. Collected fractions were diluted with 1.3 g/mL CsCl to a total volume of 11.5 mL and subject to a second round of ultracentrifuged using an SW40ti rotor for 16 hours at 10 °C. Bands were again collected by sidewall puncturing and stored in CsCl with 2 mM MgCl_2_ at 4 °C until further use. Immediately prior to use, samples were buffer exchanged and concentrated into suspension buffer (PBS with 187 mM NaCl, 2 mM MgCl_2_, 6 µM Tween 80, and 0.1 mM EDTA) using Amicon® Ultra centrifugal filter units with 100 kDa NMWCO (Millipore Sigma, UFC9100).

### Cryo-EM Data collection

Quantifoil 400 mesh holey (2 µm) grids (Electron Microscopy Sciences) were cleaned for 30s using a Fischione Instrument NanoClean. AdVLP-7 at 1 mg/ml (4 µl) were applied to the grids for 3 s, blot time of 4 s, blot force of 2, drain time of 0 s, a temperature of 4 °C, a relative humidity of 100%, and plunge frozen into a bath of liquid ethane using an FEI Mark IV. Data collection statistics are shown in Table S1. Frame alignment and dose weighting were done on-site using the MotionCor2 package with default parameters and a patch 5 (34). Initial single particle analysis: CTF estimation, manual particle picking, template picking, 2D classification, ab inito structure synthesis, heterogeneous and non-uniform refinement were performed with cryoSPARC (Fig. 1S) (35). The cryoSPARC cs and csg files were converted to Relion star files using the pyem software (36). The star and mrc files were converted to the Frealign format using in-house written scripts (37). Refinement with icosahedral symmetry (100 rounds) was performed with Frealign. Particle orientation parameters were converted to Relion using in-house written scripts. Symmetry expansion, binary mask synthesis, signal subtraction, focused classification, and 3D refinement were performed using Relion (38). Outlines in yellow (Fig. S1) and cyan define the iASU and PB masked used for focused classification, respectively. The coordinates for the AdV5 *ts1* structure were used in both instances. Initial fitting and 15 Å map synthesis were performed with UCSF Chimera (39). A cylinder mask generated with EMAN2 was used for 3D classification of the PB-Fiber (40). An AlphaFold3 model of each subunit was manually fitted into the density using UCSF Chimera (39, 41). Model building was performed with Coot, map post processing performed with Phenix *resolve_cryo_em*, and refinement performed with Phenix *real_space_refine* (42-44). Default parameters were used for the program OccuPy (19).

## Acknowledgments

KM prepared and screened cryo-EM grids. RM, HMK, MDR and KW performed virus expression and purification. KM and RK performed data collection, while RK performed data analysis and structure refinement. PG, RK, and JM designed experiments. All authors contributed to the preparation, revision, and finalization of the manuscript. Grid preparation and screening were conducted at the Imaging Facility of City University of New York (CUNY) Advanced Science Research Center (ASRC). Cryo-EM data was collected at the New York Structural Biology Center (NYSBC). We would like to acknowledge the scientific and technical assistance provided by the NYSBC and the CUNY ASRC.

