## Supplemental for "The structure of the Human Adenovirus 7 virus-like particles reveals that pentons and core-genome promote hexon-pIIIa interactions during capsid maturation"

Kiyano Madoo[1,6], Ryan Mazboudi[2,5, 6], Robert Alan Kuschner[3], Paul Gottlieb[4], Jose M. Galarza[2]
Reza Khayat[1]
[1] Department of Chemistry and Biochemistry, The City college of New York, New York, NY 10031, USA
[2] TechnoVax, 2 Hilltop Rd, Kinnelon, NJ 07405, USA
[3] Capsidvax, Arlington, VA 22201, USA
[4] CUNY School of Medicine, The City college of New York, New York, NY 10031, USA
[5] Planet Pharma, 900 Ridgebury Rd, Ridgefield, CT 06877
[6] These authors contributed equally

**This PDF file includes:**

Supporting text
Figures S1 to S5
Tables S1 to S4

**Other supporting materials for this manuscript include the following:**

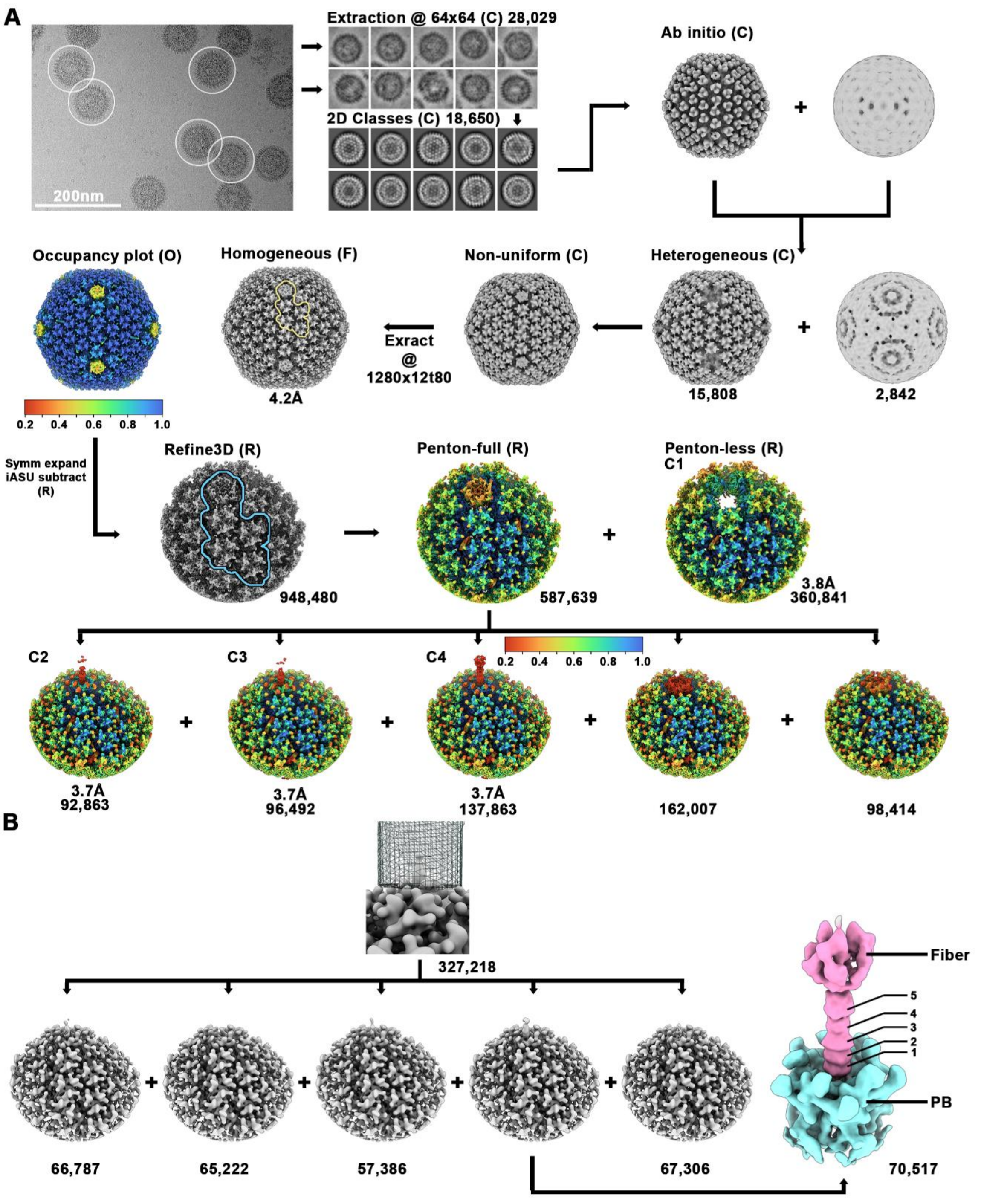

Fig. S1. **Flowchart of cryo-EM data processing.** A) Cryo-EM data processed using a combination of cryoSPARC, Frealign, and Relion. Particle picking, extraction, 2D classification, ab initio structure synthesis, heterogeneous and non-uniform refinement were performed with cryoSPARC (labeled C). Homogeneous refinement was performed with Frealign (F). Signal subtraction, focused classification, and focused refinement were performed with Relion (R). The program OccuPy (O) was used to determine the occupancy of the subunits in the Frealign and Relion maps. The yellow outline, which surrounds the iASU of the Frealign generated map, describes the region defined by a mask used for one round of signal subtraction by Relion. The one round of signal subtraction partially removed the boundary of the masked region. This is demonstrated by the reduced occupancy of the hexons in the subsequent classes. We note that this does not affect the analysis.

The blue outline surrounding the first subtracted map identifies the region modeled with atomic coordinates. The postprocessed density, generated using Phenix resolve_cryo_em, is shown in Fig. 2. B) Data processing for the PB-fiber complex. 3D classification was performed with a tau2-fudge factor of 500.

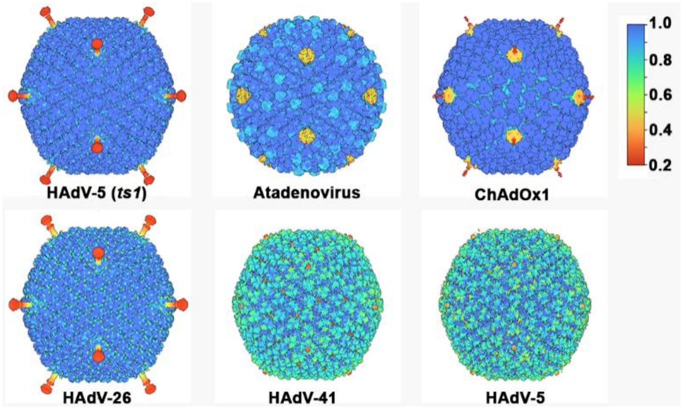

**Fig. S2. Occupancy plots.** Surface representation of AdV maps deposited in the EMDB colored according to occupancy as measured with the program OccuPy. The key shown on far right indicates the measured occupancy.

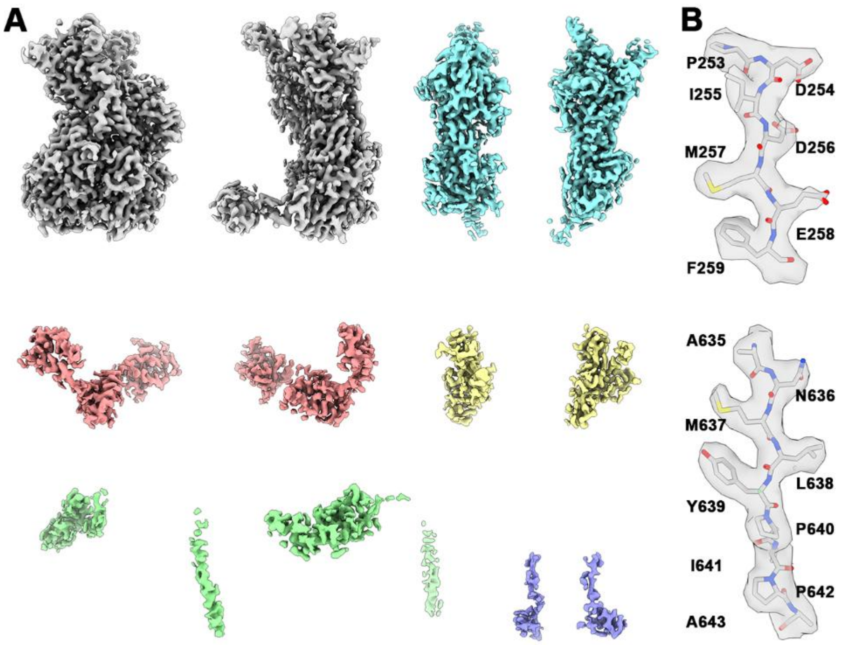

**Fig. S3. Subunit density maps.** A) Surface representation of postprocessed segmented maps used for modeling the coordinates. The subunits are colored as described in Fig. 1. The segmented maps demonstrate that side chain density is well defined and interpretable. B) Extracted density from different regions of the hexon with atomic models.

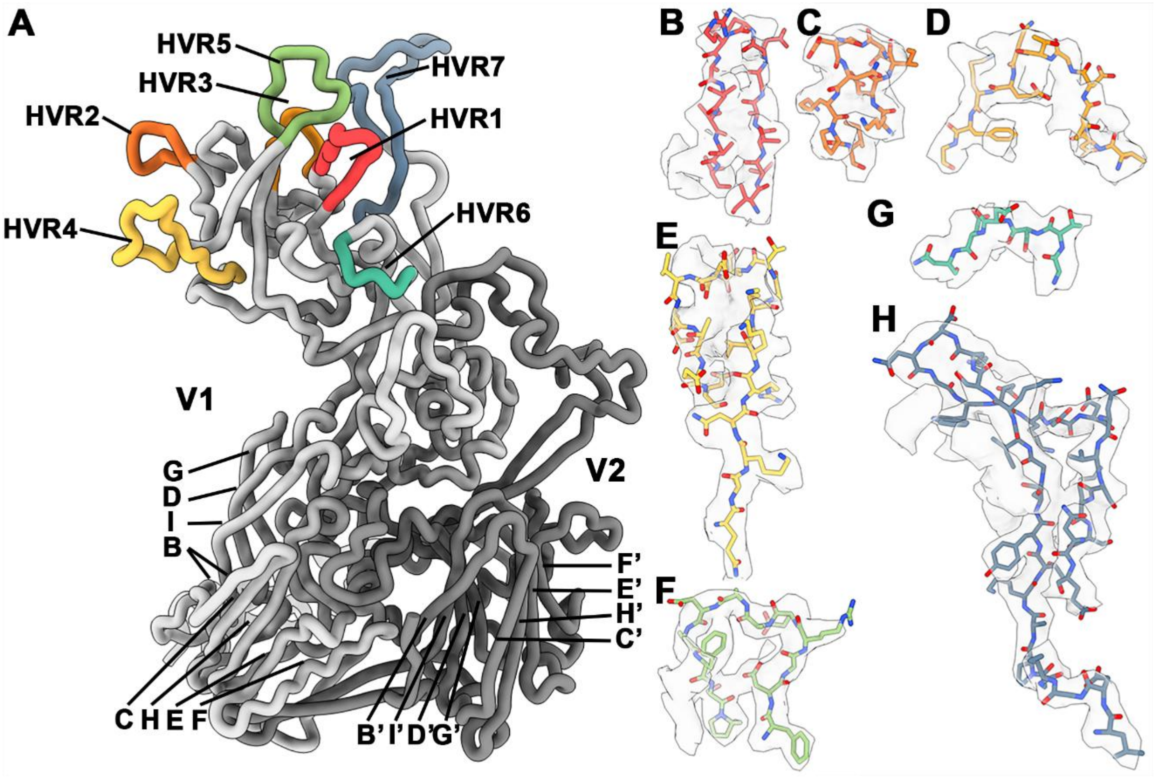

**Fig. S4. Hexon highly variable regions (HVR).** A) Licorice representation of hexon protomer with variable regions and β-strands (CHEF/BIDG) labeled. The hexon HVR are 136-148, 171-180, 198-208, 235-252, 260-271, 295-299, 406-436 (Crawford-Miksza and Schnurr JVI 1996). B-H) Electron density with modeled coordinates demonstrates that the seven HVR can be modeled.

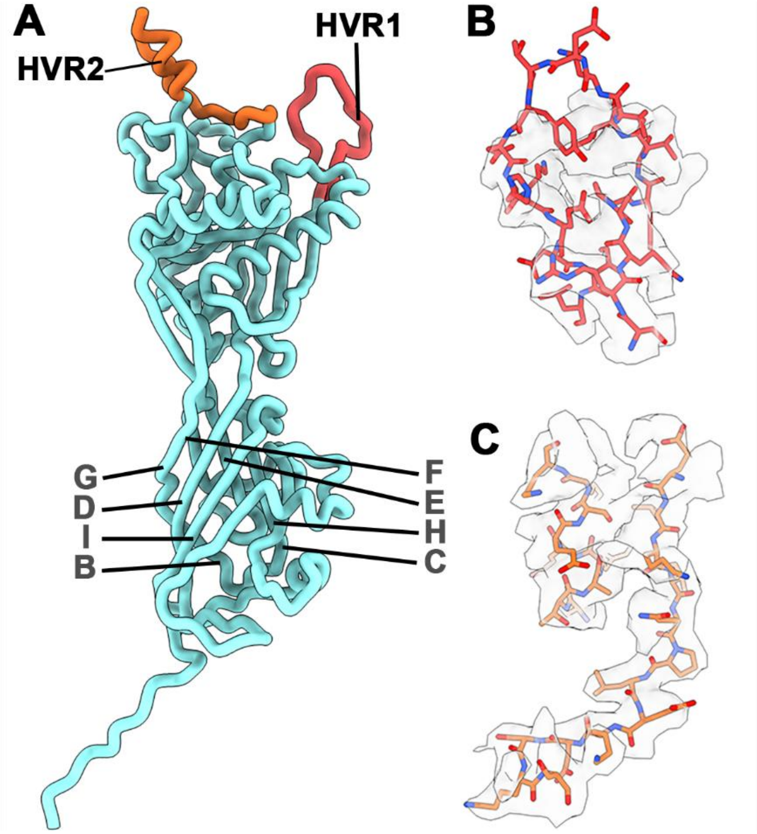

**Fig. S5. Penton highly variable regions (HVR).** A) Licorice representation of a penton protomer with variable regions and β-strands (CHEF/BIDG) labeled. The penton HVR are 150-170, 298-362. B-C) Electron density with modeled coordinates demonstrates that HVR1 can be modeled. Only a portion of HVR2 is observed**.**

**Table S1. Cryo-EM data collection, image analysis, modeling, refinement, and validation statistics**

| | VLP | C1 | C2 | C3 | C4 | PB-Fiber |
|---|---|---|---|---|---|---|
| **Data collection and processing** | | | | | | |
| Microscope | Titan Krios | Titan Krios | Titan Krios | Titan Krios | Titan Krios | Titan Krios |
| High tension (kV) | 300 | 300 | 300 | 300 | 300 | |
| Detector | Gatan K3 | Gatan K3 | Gatan K3 | Gatan K3 | Gatan K3 | Gatan K3 |
| Nominal Magnification | 81,000 | 81,000 | 81,000 | 81,000 | 81,000 | 81,000 |
| Cs (nm) | 2.7 | 2.7 | 2.7 | 2.7 | 2.7 | 2.7 |
| Data Acquisition Software | Leginon | Leginon | Leginon | Leginon | Leginon | Leginon |
| Electron dose ($e^{-}$ $Å^{-2}$) | 53 | 53 | 53 | 53 | 53 | 53 |
| Dose rate (e- $A^{-2}$ $s^{-1}$) | 21.3 | 21.3 | 21.3 | 21.3 | 21.3 | 21.3 |
| Pixel Size | 1.083 Å | 1.083 Å | 1.083 Å | 1.083 Å | 1.083 Å | 1.083 Å |
| Defocus range ($\mu$m) | -0.5 to -2.0 | -0.5 to -2.0 | -0.5 to -2.0 | -0.5 to -2.0 | -0.5 to -2.0 | -0.5 to -2.0 |
| Number of Movies | 5,608 | 5,608 | 5,608 | 5,608 | 5,608 | 5,608 |
| Number of particles | 15,808 | 360,841 | 92,863 | 96,492 | 137,863 | 70,517 |
| Symmetry imposed | I1 | C1 | C1 | C1 | C1 | C1 |
| Resolution (Å) | 4.2 | 3.8 | 3.7 | 3.9 | 3.7 | 6.1 |
| FSC threshold | 0.143 | 0.143 | 0.143 | 0.143 | 0.143 | 0.143 |
| **Refinement** | | | | | | |
| Initial model used (PDB code) | | Alpha Fold 3 | Alpha Fold 3 | Alpha Fold 3 | Alpha Fold 3 | |
| Non-hydrogen atoms | | 93,176 | 96,628 | 97,999 | 98,114 | |
| Protein residues | | 12,183 | 12,617 | 12,799 | 12,821 | |
| R.M.S. deviations | | | | | | |
| Bond lengths (Å) | | 0.005 | 0.006 | 0.006 | 0.005 | |
| Bond angles (°) | | 0.925 | 0.877 | 0.992 | 0.981 | |
| **Validation** | | | | | | |
| MolProbity score | | | | | | |
| Clashscore | | 3.0 | 3.2 | 4.2 | 4.2 | |
| EMRinger | | 3.5 | 3.0 | 3.3 | 3.7 | |
| Poor Rotamers (%) | | 0.04 | 0.02 | 0.05 | 0.05 | |
| Ramachandran (%) | | | | | | |
| Favored | | 91.7 | 90.8 | 91.4 | 91.6 | |
| Allowed | | 7.8 | 8.6 | 0.5 | 7.9 | |
| Disallowed | | 0.5 | 0.6 | 8.1 | 0.5 | |
| Fit to map (CCmask) | | 82.9 | 80.1 | 82.6 | 83.7 | |
| **Accession codes** | | | | | | |
| EMDB (maps) | 72794 | 72793 | 72774 | 72773 | 72772 | 72771 |
| PDB (model) | | 9YD0 | 9YCJ | 9YCI | 9YCH | |

**Table S2. Modeled coordinates.**

| | Hexon | Penton | pIIIa | pVIII$^{U}$ | pVIII$^{V}$ | IX | pVI |
|---|---|---|---|---|---|---|---|
| C1 | A 4-935<br>B 5-931<br>C 8-936<br>D 3-934<br>E 6-934<br>F 5-935<br>G 5-936<br>H 4-934<br>I 6-935<br>J 5-935<br>K 5-936<br>L 4-935 | None | 28-104 | 2-61;<br>181-226 | 2-61;<br>181-227 | P: 8-70;<br>97-132<br>Q: 8-70;<br>92-125<br>R: 8-70;<br>100-134<br>S: 8-69;<br>99-134 | W: 4-19; 23-31<br>Y: 4-20; 24-31<br>Z: 5-19, 23-34<br>0: 5-13, 27-33<br>1: 5-31<br>2: 5-31<br>3: 5-19; 25-33<br>4: 5-19; 22-33 |
| C2 | A 2-935<br>B 5-935<br>C 3-935<br>D 3-934<br>E 5-934<br>F 5-936<br>G 5-935<br>H 3-935<br>I 5-935<br>J 5-935<br>K 5-937<br>L 3-935 | 52-306,<br>350-543 | 28-108 | 2-61;<br>181-226 | 2-61;<br>181-227 | P: 8-70;<br>97-132<br>Q: 8-70;<br>92-125<br>R: 8-70;<br>100-134<br>S: 8-69;<br>99-134 | W: 4-31<br>Y: 4-20; 24-31<br>Z: 5-19, 27-30<br>0: 5-13, 27-33<br>1: 5-19; 24-31<br>2: 5-19; 24-32<br>3: 5-19; 29-30<br>4: 5-19; 22-31 |
| C3 | A 2-935<br>B 5-936<br>C 3-937<br>D 3-934<br>E 6-934<br>F 5-937<br>G 5-936<br>H 4-935<br>I 5-935<br>J 5-935<br>K 5-937<br>L 3-935 | 38-306,<br>350-543 | 28-218,<br>226-266 | 2-61;<br>181-226 | 2-61;<br>181-227 | P: 8-70;<br>97-132<br>Q: 8-70;<br>92-125<br>R: 8-70;<br>100-134<br>S: 8-69;<br>99-134 | W: 4-31<br>Y: 4-20; 24-31<br>Z: 5-19, 23-34<br>0: 5-13, 27-33<br>1: 5-19; 24-40<br>2: 5-19; 24-32<br>3: 5-19; 29-31<br>4: 5-19; 22-30 |
| C4 | A 2-937<br>B 5-937<br>C 3-934<br>D 3-934<br>E 6-937<br>F 5-936<br>G 5-935<br>H 4-935<br>I 5-935<br>J 5-935<br>K 5-937<br>L 3-935 | 38-306,<br>350-543 | 28-218,<br>226-266 | 2-61;<br>181-226 | 2-61;<br>181-227 | P: 8-70;<br>97-132<br>Q: 8-70;<br>92-125<br>R: 8-70;<br>100-134<br>S: 8-69;<br>99-134 | W: 4-31<br>Y: 4-20; 24-31<br>Z: 4-34<br>0: 5-13, 27-33<br>1: 5-19; 25-33<br>2: 5-19; 24-32<br>3: 5-19; 25-33<br>4: 5-19; 22-33 |

**Table S3. Subunit-subunit interface (buried surface area)**

| | C1 | C2 | C3 | C4 | AdV5 *ts1* | AdV5 |
|---|---|---|---|---|---|---|
| Hex-Pent | N/A | 900 | 2700 | 3100 | 3500 | 4100 |
| Hex-pIIIa | 2100 | 2100 | 2900 | 3000 | 6100 | 5200 |
| Hex-pVIII$^{U}$ | 7900 | 8200 | 8300 | 8800 | 12600 | 12200 |
| Hex-pVIII$^{V}$ | 7600 | 8400 | 8500 | 9100 | 12200 | 13000 |
| Hex-IX | 19600 | 19900 | 19700 | 1990 | 25400 | 25300 |
| Pen-pIIIa | N/A | 0 | 1500 | 1500 | 2000 | 1900 |
| pIIIa-pVIII | 0 | 0 | 1400 | 1300 | 2900 | 2800 |
| Pent$^{Total}$ | N/A | 900 | 4200 | 4600 | 5400 | 6000 |
| pIIIa$^{Total}$ | 2000 | 2100 | 5700 | 5700 | 11000 | 9800 |
| pVIII$^{Total}$ | 15500 | 16600 | 18200 | 19200 | 27600 | 27900 |

Units are in Å$^2$

**Table S4. Table S4: Residues contributing to hexon-penton interface.**

| **Penton region** | **Interacting hexon residues** |
|---|---|
| 74-80 | C2: 682-683<br>C3: 78, 678, 682-683<br>C4: 78, 85, 351, 675, 678, 682 |
| 97-109 | C2: 331, 351<br>C3: 348-352, 637, 653, 654, 931-935<br>C4: 350-352, 637, 653-655, 931-934 |
| 426-429 | C2: 74, 76<br>C3: 72, 74, 81, 83, 336, 343, 567<br>C4: 72, 74, 81, 83, 336, 340, 343, 567 |
| **Hexon region** | **Interacting penton residues** |
| 75-78 | C2: 397-400, 425, 427-429<br>C3: 85, 396-399, 429-430<br>C4: 85, 397-399, 427, 429, 430 |
| 331-334 | C2: 81, 82, 84, 85<br>C3: 82, 92, 397, 398, 428, 509, 511, 513<br>C4: 82, 92, 396-398, 428, 509, 511, 513 |
| 340-345 | C2: N/A<br>C3: 425,426,428,505<br>C4: 425,426,428,505 |
| 676-679 | C2: 81, 82, 84, 85<br>C3: 81, 82, 84<br>C4: 82, 84, 101, 108, 110, 425, 428, 505, 513 |
| 927-936 | C2: N/A<br>C3: 58, 109, 110, 112, 399, 400, 507<br>C4: 56, 58, 59, 105, 110, 112 |
